# Reexamining the essentiality of Pdi1 in yeast – A *PDI1* knockout is viable in *Komagataella phaffii* and still produces recombinant disulfide bonded proteins

**DOI:** 10.1101/2024.08.21.609038

**Authors:** Arianna Palma, Viktoria Kowarz, Brigitte Gasser

**Author notes:** Corresponding author: Brigitte Gasser, Institute of Microbiology and Microbial Biotechnology, Department of Biotechnology, BOKU University, Muthgasse 18, Vienna, Austria.

## Abstract

Protein disulfide isomerase 1 (Pdi1) and Ero1 form the main oxidative folding axis in the endoplasmic reticulum (ER). Despite having additional PDI family members, Pdi1 has long ago been determined to be essential in baker’s yeast and this concept was inferred to other yeast species. In this study, we reexamine the gene essentiality of Pdi1 in the methylotrophic yeast *Komagataella phaffii*. Strikingly, the absence of Pdi1 does not cause lethality, but even allows for folding and secretion of heterologous proteins in a homogeneous redox state. Remarkably, while sensitivity to externally added folding stressors, such as tunicamycin and DTT is increased in *pdi1*Δ knock-out cells, in non-stressed growth conditions they do not display upregulation of folding stress markers, such as the master chaperone Kar2 or the unfolded protein response regulator Hac1, suggesting homeostatic adaptation.

## Introduction

Human PDI (hPDI) and its homolog Pdi1 in the yeast *Saccharomyces cerevisiae* are by far the most studied PDI family members both *in vitro* and *in vivo*, and thought to be the main catalyst for disulfide formation and isomerization (1, 2). Several PDI family members have been identified, of which PDI (or Pdi1p in yeast) possess both disulfide oxidase as well as isomerase activity and is the most researched representative. The *Homo sapiens* species counts more than twenty PDI family proteins, whose functional properties and substrate specificities have been only partially elucidated. Baker’s yeast *S. cerevisiae* presents five PDI family members (Pdi1p, Mpd1p, Mpd2p, Eug1p, Eps1p), whose specific roles have also not been entirely deciphered (1–3).

Pdi1p is essential in *S. cerevisiae*, where *pdi1*Δ mutants were shown to be lethal (4–6), although the single overexpression of the other four PDI family members was proven to suppress lethality (7–11). Human P4HB knockout cell lines exist and are commercialized, as well as conditional knockout mice models, whereas non-conditional knockout mice are embryonically lethal (12, 13). Isomerization seems to be more crucial for homeostasis of protein folding in higher eukaryotic cells, which express proteins with more intricate disulfide patterns, while only 6% activity seems to be sufficient for sustaining the growth of yeast cells (4, 14). Although oxidation only seems to be sufficient to rescue growth in *S. cerevisiae*, the oxidative folding process is greatly compromised by the lack of isomerase activity (14, 15). Overall, the proof-reading activity over non-native disulfide bonds is suggested to be the distinguishing function of Pdi1, which differentiates it from its numerous *cis*- and *trans*-species relatives (4, 15, 16).

In our recent work, we characterized the kinetics of Pdi1 from the methylotrophic yeast *Komagataella phaffii* (syn *Pichia pastoris*), a popular host for the secretion of industrial and biopharmaceutical protein targets (17). We showed that KpPdi1 can perform faster oxidation of disulfide bonds comparing to human PDI, but less efficient isomerization. Furthermore, we isolated and characterized a novel PDI family member, Erp41, which is shared almost exclusively among few non-conventional fungi but does not exist in *S. cerevisiae* (18). Erp41 revealed to be an oxidase with unprecedented catalytic activity in partnership with glutathione, while being a less efficient isomerase comparing to KpPdi1. Given the presence of two strong oxidases in the ER of *K. phaffii*, we investigated whether Pdi1 is truly essential in this yeast.

## Results

### Generation of a *K. phaffii pdi1Δ* strain and its phenotypical characterization

We attempted knocking out the *PDI1* gene of *K. phaffii* by CRISPR-Cas9 homology directed recombination. Transformation plates revealed a majority of average-sized colonies and a minority of small-sized colonies, which were confirmed to correspond to full knock-outs (Fig. 1, Fig. *S*1).

**Figure 1.**
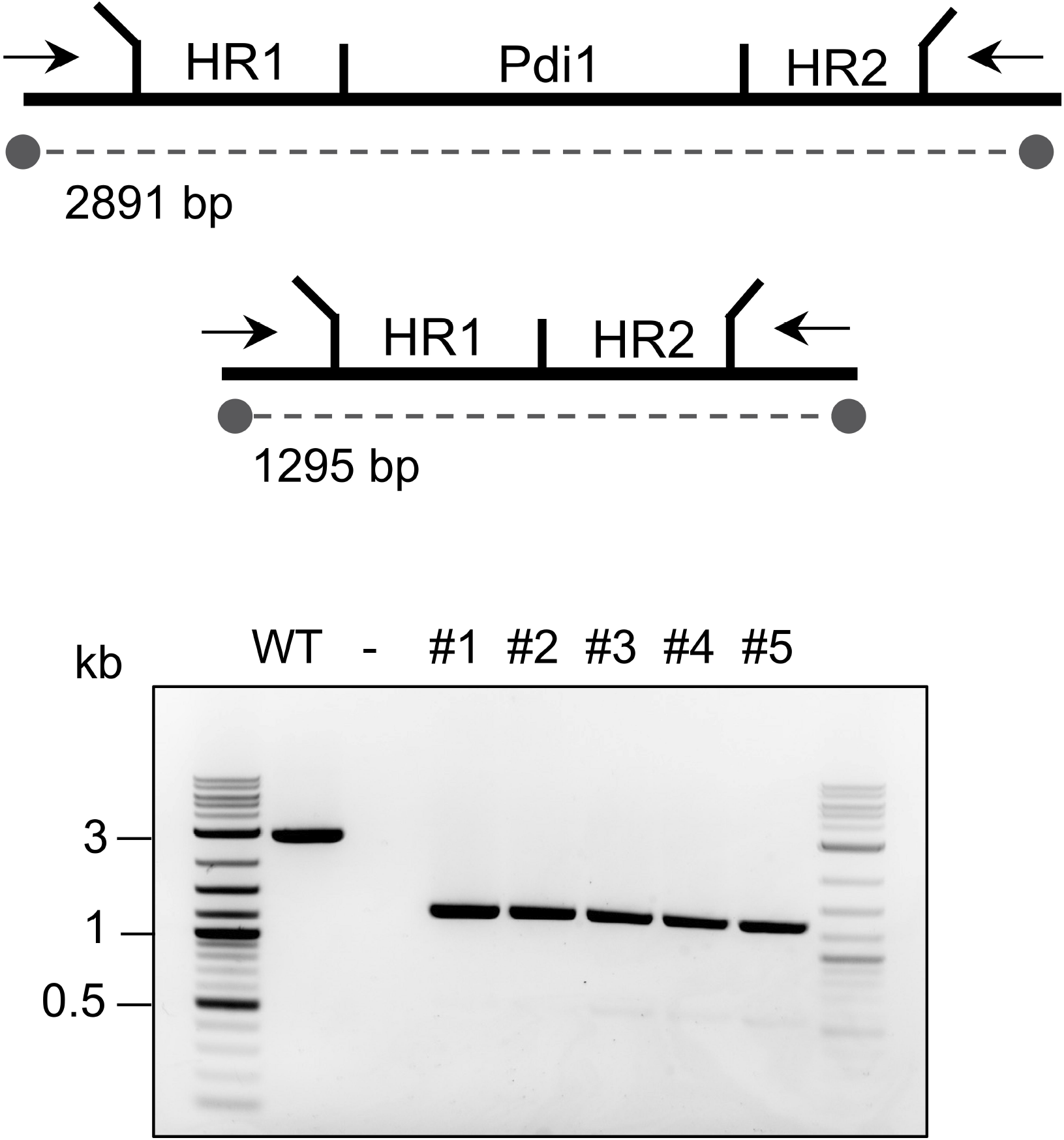
Confirmation of *K. phaffii* Pdi1 knock-out by locus PCR. *A*. Schematic comparison of the wild-type (WT) and knock-out (#1-5) loci with agarose gel displaying locus PCR results. Complementary gels showing the absence of the *PDI1* gene are provided in Fig. *S*1.

Differences in viability between wild-type and *pdi1*Δ clones were investigated though growth assessment on solid and liquid media, on both rich and minimal (YPD and YNBGlc). In both conditions, the *pdi1*Δ clones showed consistently slower growth comparing to the wild-type reference (Fig. 2). Despite slower growth, the morphology of the spots appeared normal and by itself did not reveal underlying stress (Fig. 2*A*). Cultures were then supplemented with stressors and folding interferants reported to upregulate the unfolded protein response (UPR), such as dithiothreitol (DTT), a strong reductant, diamide, an oxidant, and tunicamycin, which impairs glycosylation (14, 19). These stress-inducing molecules further affected growth of the knock-out strains, especially when combined with media lacking rich components. Sensitivity of *pdi1*Δ to DTT and tunicamycin was clearly increased, while diamine did not show an effect compared to the untreated control (Fig. 2A). Analysis of growth profiles in liquid YNBGlc, revealed that the *pdi1*Δ clones duplicate at approximately half the rate of the wild-type reference, having doubling times of 0.04 and 0.1 OD/h, respectively (Fig. 2B). Supplementation of liquid YNB with tunicamycin mildly affected the wild-type reference (growth rate was reduced to 70%) and drastically affected the growth of the *pdi1*Δ mutant, which dropped to approximately 15% of the wild-type. However, none of the assayed conditions was effective in suppressing viability.

**Figure 2.**
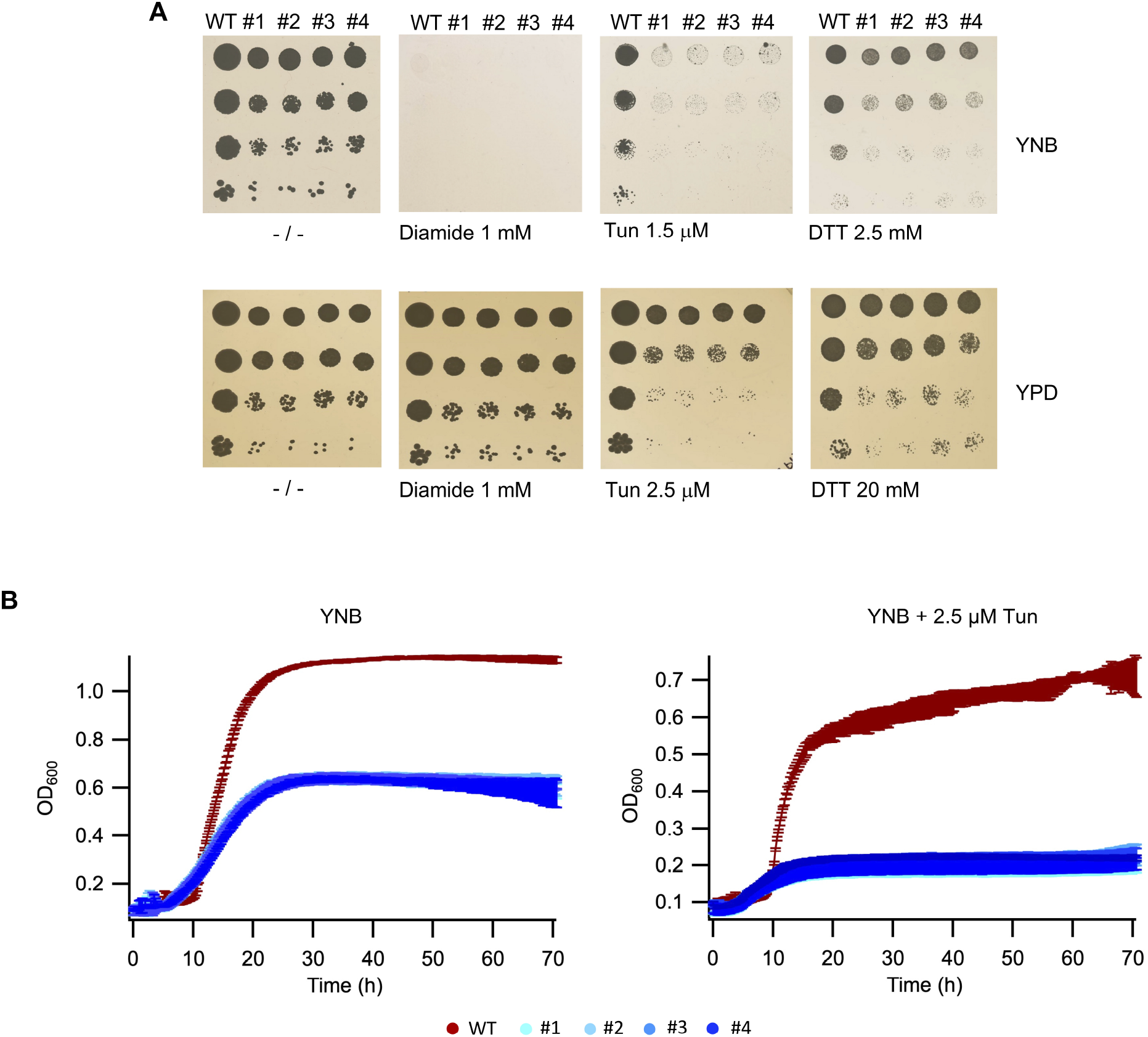
Phenotypical characterization of the Pdi1 knock-out strains. *A*. Spotting assays of wild-type (WT) and KO strains (#1-4) performed on YPD and YNBGlc (containing 2% glucose), plain or supplemented with several concentrations of tunicamycin, diamide or DTT. Results obtained with 1 mM diamide, 1.5 µM tunicamycin, 2.5 mM DTT in YNB, 2.5 µM tunicamycin, and 20 mM DTT in YPD were selected for display. *B*. Growth curves of wild-type (WT) and KO strains (#1-4) in liquid YNBGlc, plain and supplemented with 2.5 µM tunicamycin.

### Folding competence of *K. phaffii pdi1*Δ towards non-native model proteins

Given that the absence of the Pdi1 allowed *K. phaffii* to be vital (albeit showing reduced growth), in contrast to previous studies conducted on *S. cerevisiae*, we analyzed the ability of the *pdi1*Δ clones to fold heterologous disulfide-containing models. For this purpose, we expressed porcine trypsinogen (Trp), which contains six disulfides, and a single-chain variable fragment (scFvM) with two disulfides. Both model proteins were produced and secreted by the *pdi1*Δ clones (Fig. 3). Clean secretion was observed, as there were no additional bands appearing in the *pdi1*Δ strains compared to the wild-type. Nevertheless, when normalized the titers to the wet cell weight (WCW) at harvest, for both reporter proteins the biomass specific product yields (titer/WCW) were drastically lowered in *pdi1*Δ compared to the wild-type strain. Specifically, yields for scFvM were 40% of the wild-type, while trypsinogen production was reduced to 27% of the WT.

**Figure 3.**
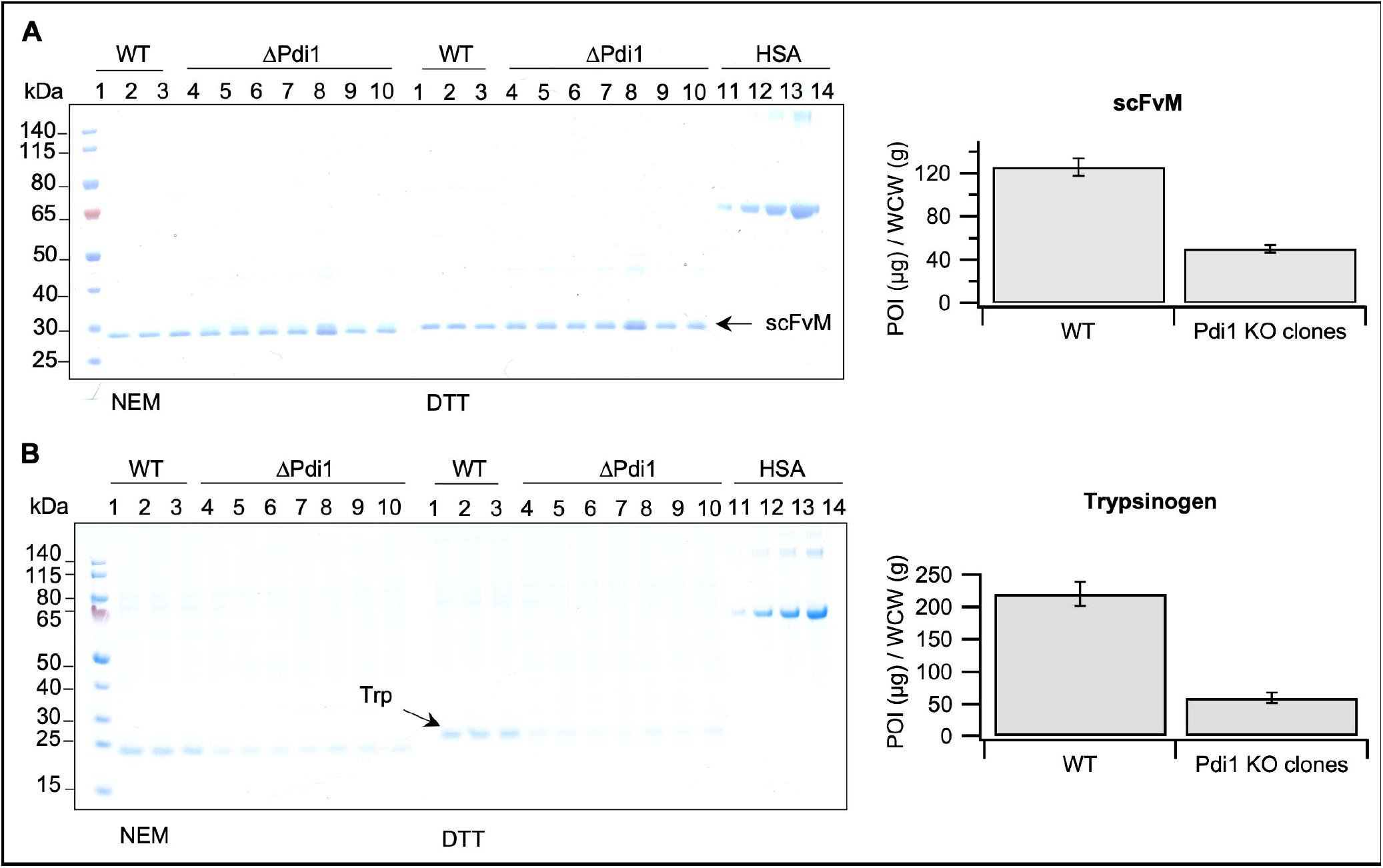
Secretion profiles of recombinant model proteins in *pdi1*Δ. *A*. scFvM and *B*. porcine trypsinogen. SDS-PAGE (NEM-blocked and DTT-reduced) profiles and secretion titers expressed in µg of secreted protein per gram of wet cells. SDS-PAGE gels display 4 human serum albumin (HSA) standards at 0.1, 0.3, 0.6 and 1 µg.

**Figure 4.**
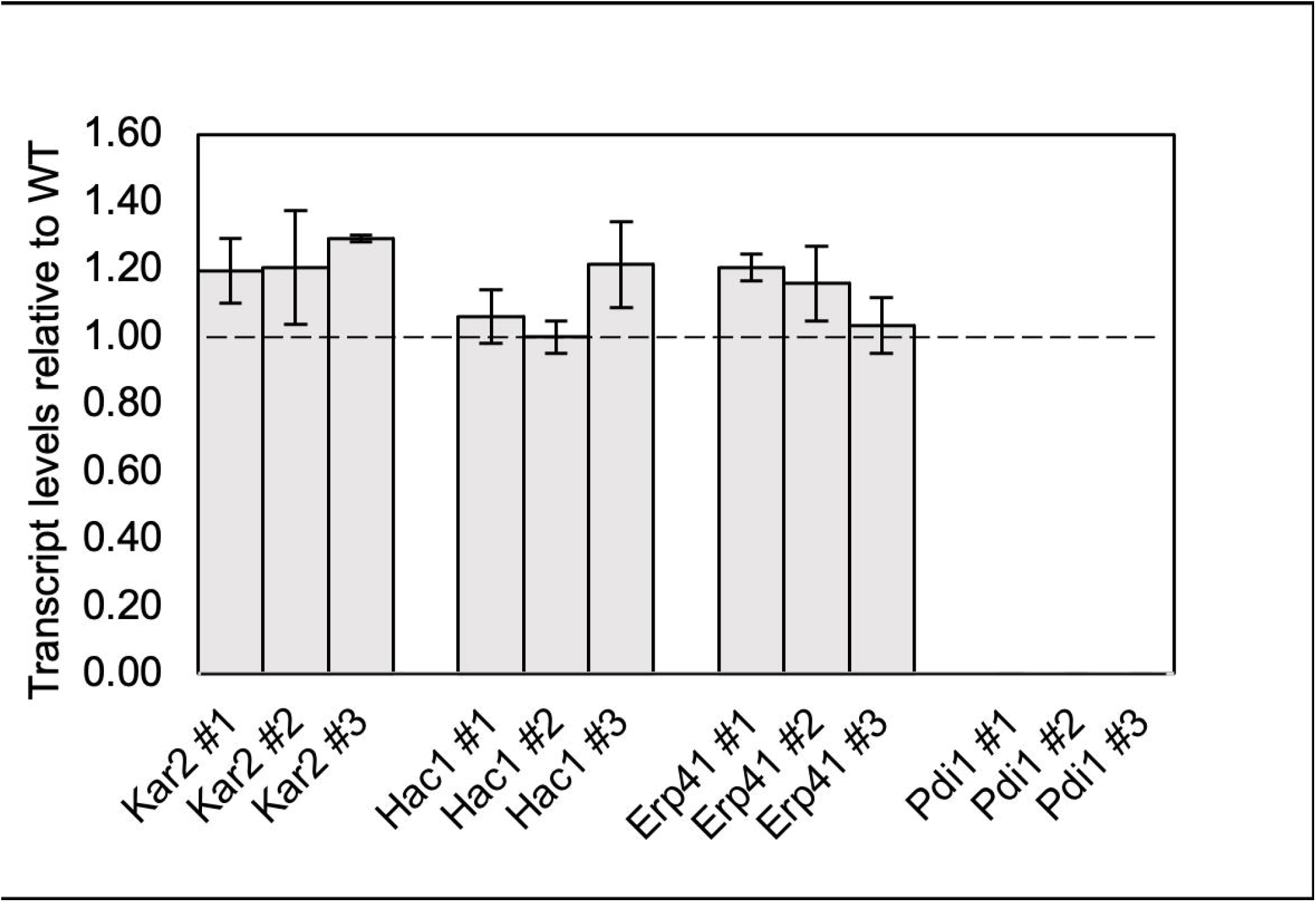
Transcript levels of UPR markers. Transcript levels relative to WT values for *KAR2, HAC1, ERP41* and *PDI1* (negative control) are reported. Data is presented as mean and standard deviation of at least three technical replicates per clone.

Despite the absence of Pdi1, *K. phaffii* was still able to produce and secrete recombinant disulfide-containing proteins. Thus, we set out to investigate the disulfide arrangement of both model proteins with mass spectrometry. Specifically, two *pdi1*Δ samples and one wild-type reference were treated with NEM to label free cysteines and analyzed through electrospray ionization MS. All knock-out samples displayed comparable chromatographic profiles with respect to the wild-type, with one main peak corresponding to the mass of the oxidized targets obtained, followed by two minor peaks generated by aspecific NEM adducts (NEM bound to oxidized protein target) (Table1). Therefore, both wild-type and *pdi1*Δ mutant strains delivered fully folded trypsinogen and scFvM, however the latter delivered significantly lower titers.

### Transcript level analysis of UPR markers

As the *K. phaffii pdi1*Δ knock-out mutants seemed to be viable and functional, we proceeded to test whether this was due to the upregulation of genes related to the UPR pathway, which could compensate for the absence of Pdi1 both for folding the native and recombinant disulfide-bonded proteins. The most commonly assayed genes to verify whether the UPR is upregulated are *HAC1*, encoding the main transcription factor which initiates the UPR response, and *KAR2*, a very abundant ER chaperone, which acts as a master regulator for the ER folding homeostasis (20–22). Surprisingly, neither *HAC1* nor *KAR2* were significantly upregulated in the *pdi1*Δ clones, suggesting that the strains adapted to the absence of the Pdi1 and tuned their homeostasis accordingly, without the constitutive intervention of stress responses. Additionally, we checked the transcript levels of the *ERP41* gene, which acts an efficient oxidant and might be induced to take over Pdi1 functionalities. Also in this case, no significant upregulation could be observed.

**Table 1:** Electrospray ionization mass spectrometry analysis of scFvM and porcine trypsinogen. One supernatant sample purified from the wild-type (WT) strain and two from *pdi1*Δ were analyzed. Both theoretical and experimental masses are reported as average masses. Theoretical average masses refer to the reduced state of the proteins unless marked otherwise.

| Protein | Theoretical Mass (Da) | Sample | Experimental Mass (Da) | ΔMass (Da) | # Disulfides (Da) |
| --- | --- | --- | --- | --- | --- |
| scFvM | 27700.44 | WT | 27696.7 | -3.74 | 2 |
|  |  |  | 27821.9 | +121.46 (1 NEM) |  |
|  |  |  | 27947.2 | +246.76 (2 NEM) |  |
|  |  | KO #1 | 27696.8 | -3.64 | 2 |
|  |  |  | 27822.2 | +121.76 (1 NEM) |  |
|  |  |  | 27947.3 | +246.86 (2 NEM) |  |
|  |  | KO #2 | 27696.8 | -3.64 | 2 |
|  |  |  | 27821.8 | 121.36 (1 NEM) |  |
|  |  |  | 27947 | 246.56 (2 NEM) |  |
| Trypsinogen | 24312.36 | WT | 24300.88 | -11.48 | 6 |
|  |  |  | 24425.83 | 113.47 (1 NEM) |  |
|  |  |  | 24550.86 | 238.5 (2 NEM) |  |
|  |  | KO #1 | 24300.77 | -11.59 | 6 |
|  |  |  | 24425.76 | 113.4 (1 NEM) |  |
|  |  |  | 24550.94 | 238.58 (2 NEM) |  |
|  |  | KO #2 | 24300.82 | -11.54 | 6 |
|  |  |  | 24425.5 | 113.14 (1 NEM) |  |
|  |  |  | 24550.59 | 238.23 (2 NEM) |  |

## Discussion

The long-determined gene essentiality of Pdi1 in *S. cerevisiae* has been attributed to its oxidase and isomerase function, the latter being less prominent in its paralogs (4, 5, 16). In this study, we discovered that Pdi1 is not essential for viability in the non-conventional yeast *K. phaffii*. In fact, the Pdi1 knock-out strains that we generated survive even under chemically induced redox and folding stress, with the only observable phenotypical trait being slower growth rates comparing to the wild-type reference.

Remarkably, the Pdi1 deletion does not seem to affect the redox homogeneity of two disulfide-rich heterologous protein targets, scFvM and porcine trypsinogen, while causing lower titers. A precondition for a protein to successfully transit through the secretory pathway is to be in a folding state that, if not native, must not cause major aggregation or insolubility. We observed that 100% of the secreted target matched the correct molecular weight corresponding to the oxidized protein, largely suggesting that the cystine patterns were homogeneous. This might imply that neither the Pdi1 oxidation activity neither its isomerization/proofreading activity are strictly necessary to deliver correctly folded non-native protein targets, although its presence increases production and secretion rates.

Furthermore, *pdi1*Δ knock-out cells did not display any sign of upregulation of UPR target genes in standard growth conditions. This is in stark contrast to previous studies analyzing the unfolded protein response in isomerase-deficient strains (expressing only the *a’* domain of Pdi1) in *S. cerevisiae* (14). Therefore, we hypothesize that *K. phaffii* adapted and retuned its homeostasis in the absence of Pdi1, possibly by slowing down translation, growth and thus secretion rates. We additionally checked the transcript levels of Erp41, the novel Pdi1 homolog we have recently characterized. Our *in vitro* studies (18) showed that Erp41 is an excellent oxidase in partnership with Ero1, and its speed is so far unmatched when associated to the GSH/GSSG redox couple. Differently from Pdi1, the *erp41*Δ knock out did not cause any observable phenotype, which in combination with the scarce presence of the Erp41 in other organisms, did not allow to identify the specific functions or folding pathways which might be covered by this unique enzyme. Theserefore, based on transcript data, the viability of *K. phaffii pdi1*Δ is not due to increased *ERP41* expression levels.

Concludingly, this work proposes to reexamine the effective role of Pdi1 in yeasts, opening questions on the extent at which reshuffling of disulfides is needed for native yeast proteins, and possibly shifting the focus to other members of the PDI family.

## Experimental Procedures

### Vector assembly

Yeast vectors were generated using Golden Gate cloning in the GoldenPiCS library adapted for *K. phaffii* (23). The list of constructs used in this study is reported in Table *S*1.

### Pdi1 knock-out generation

The Pdi1 knock-out was generated following the protocol described in Gassler *et al*. (24) for CRISPR homology-directed gene targeting in *K. phaffii*. Briefly, *K. phaffii* CBS7435 cells were transformed by electroporation with a plasmid expressing the SpCas9 gene under P_LAT1_ and the sgRNA under P_GAP_ and a linear repair template. P_LAT1_ and P_GAP_ are respectively a moderate and a strong promoter in the presence of glucose as a carbon source. To increase efficiency, two different guides targeting the 5’ of the gene were designed, one annealing to the forward strand and the other to the reverse strand. Two constructs were assembled with either nourseothricin (NTC) or G418 resistances, each carrying one guide, and transformed together. The homology regions to generate the repair template were selected 500 bp upstream and downstream of the gene, connected with a BsaI site, cloned into the Golden Gate vector system and from there amplified to obtain a linear 1000 bp fragment. Approximately 100 ng of each plasmid and 3000 ng of repair template were combined in the transformation mix. Transformants growing on NTC and G418 were screened with colony PCR, both with primers annealing outside of the targeted locus and of the region used for homology-based repair (Figure 1), and with gene-specific primers, in order to exclude reintegration events (as shown in Fig. *S*1). The list of oligonucleotides used in this study is reported in Table *S*2.

### Viability assessments

Pdi1 knock-out clones were tested for viability on solid and liquid media, respectively, through spotting assays and growth experiments with real-time OD_600_ monitoring in a Sunrise photometer (Tecan AG). Liquid precultures were carried out in 24 deep-well plates, in 2 mL YPD at 25° C and 250 rpm shaking.

For spotting assays, OD_600_ was measured after 24 h, cells were washed in 20 mM sodium phosphate, 150 mM NaCl, pH 7.0 and growth was levelled to OD_600_ 0.3 for all cultures. Serial factor 10 dilutions were made and 5 µL were spotted on YPD and YNB each with 2% glucose, either plain or supplemented with tunicamycin, diamide or DTT in different concentrations. Plates were incubated at 30°C and growth was monitored daily. Spotting plates displayed one wild-type control and four biological replicates of the Pdi1 knock-out. Three to five replicates were performed for each condition.

For growth in liquid media, preculture OD_600_ was measured after 24 h, cells were washed in YPD with and without supplemented 2.5 µM tunicamycin and levelled to OD_600_ 0.12. A volume of 180 µL was aliquoted in the wells of a transparent 96-well plate for each culture. The outermost rectangle was filled with 200 µL water, to prevent evaporation. A total of four biological replicates were tested, together with the wild type, each assayed in five technical replicates. The plate was incubated in the Sunrise photometer at 30°C, 250 rpm shaking and OD_600_ monitoring every 15 min for 72 h.

### Heterologous protein secretion screening

CBS7435 wild-type and *pdi1*Δ were transformed with integrative plasmids targeting the *AOX1* locus, expressing either porcine trypsinogen or scFvM under control of the glycolytic P_GAP_ promoter and using the *S. cerevisiae* alfa-mating factor pre-pro leader for secretion (25). scFvM was fused to a C-terminal GGS-His6 tag. The full sequence of both model proteins is provided in Table *S*3. Small-scale screenings were performed in a 24-deep-well plate (DWP) format, in which three clones of the producing WT and 8 clones of the *pdi1*Δ mutant were cultivated. Precultures were inoculated from freshly streaked colonies in 2 mL YP media supplemented with zeocin and incubated at 25°C with 280 rpm shaking overnight. Main cultures were inoculated at the starting OD_600_ of 5.0 from pelleted precultures resuspended in 1 mL ASM media supplemented with 10% soy peptone and 5% yeast extract, in glucose-limiting conditions (26). Glucose limitation was obtained by supplementing the AMS media with 50 g/L polysaccharide (EnPresso® Y Defined). A steady glucose-release rate of approx. 0.7 mg*/gh was established by the addition of 0.4% EnPresso® glucose-releasing enzyme. After incubation at 25°C at 280 for 48 h, 1 mL of each sample was pelleted, supernatant was collected, and WCW measured.

### SDS-PAGE and sample preparation

Reduced SDS-PAGE samples were prepared by mixing protein samples with loading buffer containing 100 mM DTT, left incubating for 15 min at room temperature (RT) and heated up at 95° C for 5 min. Non-reduced samples were treated with NEM at the final concentration of 25 mM for 15 min, mixed with loading buffer without reducing agent and heated up at 95°C for 5 min. Samples were run on commercial 20% acrylamide gels (Biorad) in a tris/glycine/SDS running buffer (Biorad) for 30 min at 200 mV.

### Microcapillary electrophoresis

The ‘LabChip GX/GXII System’ (PerkinElmer) was employed for quantitative analysis of secreted protein titers (trypsinogen and scFvM) in culture supernatants. The Protein Express Lab Chip (760499, PerkinElmer) was primed according to the manufacturer’s instructions with the Protein Express Reagent Kit (CLS960008, PerkinElmer). Briefly, 6 μL of culture supernatant were mixed with 21 μL of non-reducing sample buffer, heated up at 100°C for 5 min, briefly centrifuged and further mixed with 105 μL milliQ water. Samples were then centrifuged at 1200 g for 2 min and loaded the instrument. Internal standards enabled for approximate allocations to size in kDa and to determine concentrations from the detected signals. Concentration of the secreted recombinant protein were related to the WCW to determine the product yield (µg product per gram of WCW).

### Purification of protein samples

Prior to MS analysis, samples were either purified or buffer exchanged. 1 mL trypsinogen supernatant samples were buffer exchanged with 10 kDa Amicon filters in 10 mM phosphate buffer at pH 7.3. 1 mL scFvM supernatants were pre-equilibrated at pH 7.3 with phosphate buffer and purified on Poly-Prep columns (Biorad) with 0.4 mL bed volume of HisPur Cobalt Agarose resin (ThermoFischer Scientific), as described by Gaciarz *et al*. (27), with a final elution step in 50 mM EDTA in phosphate buffer at pH 7.3. Next, samples were diluted to 0.1 mg/ml in 10 mM sodium phosphate pH 7.3 and incubated for 10 min in 25 mM NEM.

### Intact protein mass spectrometry

4 µL of the protein solution were directly injected into an LC-ESI-MS system. A gradient ranging from 15 to 80% acetonitrile in 0.1% formic acid, using a Waters BioResolve column (2.1 x 5 mm), was applied at a flow rate of 400 µL/min over a 9-minute gradient duration. Detection utilized a TOF instrument (Agilent 1290 Infinity II UPLC coupled to Agilent Series 6230B TOF) equipped with the Jetstream ESI source in positive ion. Calibration of the instrument was conducted using an ESI calibration mixture from Agilent. Data processing was performed using MassHunter BioConfirm B.08.00 (Agilent), and spectral deconvolution was carried out using MaxEnt.

### Quantitative PCR

Liquid cultures for transcript level determination were performed in 10 mL YNB media + 2% glucose, inoculated at a starting OD_600_ of 0.5, at 30°C and 180 rpm shaking. After 14 h, when cells were still actively growing, 1 mL of sample was collected from shaking flasks and RNA was extracted with TRI reagent according to the manufacturer’s instructions (Ambion, USA). To remove residual DNA, the RNA samples were treated with the DNA-free™-kit (Ambion), according to the manufacturers’ manual. Subsequently, RNA quality, purity and concentration were analysed by gel electrophoresis as well as spectrophotometric analysis using a NanoDrop 2000 (Thermo Scientific). Synthesis of cDNA was done with the Biozym cDNA Synthesis Kit according to the manufacturer’s manual. For qPCR analysis appropriate amounts of the cDNA (for transcript level analysis) or genomic DNA (for gene copy number analysis) were mixed with water, primers and 2x qPCR S’Green BlueMix (Biozym Blue S’Green qPCR Kit) according to the supplier’s recommendations and measured in a real-time PCR cycler (Rotor-Gene, Qiagen). All samples were measured at least in technical triplicates. Actin was measured as a reference constitutive gene. Data analysis was performed with the Rotor-Gene software employing the Comparative Quantitation (QC) method, using *ACT1* as reference gene. All primers used for qPCR are reported in Table *S*2.

## Acknowledgments

This work was funded by the Marie Skłodowska-Curie Actions Innovative Training Network of the European Union’s Horizon 2020 Program under grant agreement no. 813979 (SECRETERS). Further support was obtained by the Austrian Federal Ministry of Labour and Economy (BMAW), the Austrian Federal Ministry of Climate Action, Environment, Energy, Mobility, Innovation and Technology (BMK), the Styrian Business Promotion Agency SFG, the Standortagentur Tirol, the Government of Lower Austria, the Business Agency Vienna and BOKU through the COMET Funding Program managed by the Austrian Research Promotion Agency FFG. The use of the facilities of the BOKU Mass Spectormetry core facility is gratefully acknowledged. We thank dearly Dr. Clemens Grünwald-Gruber and Mr. Daniel Maresch for performing the mass spectrometry measurements.

## Data availability

All data is included in the paper.

