## Supplementary Material for "Reexamining the essentiality of Pdi1 in yeast – A *PDI1* knockout is viable in *Komagataella phaffii* and still produces recombinant disulfide bonded proteins"


**Figure S1. Pdi1 knock out confirmation by locus PCR**. *A*. Schematic comparison of the wild-type and knock out loci, primer design and theoretical amplicon sizes. *B*. Agarose gel displaying locus and gene-specific PCR results.

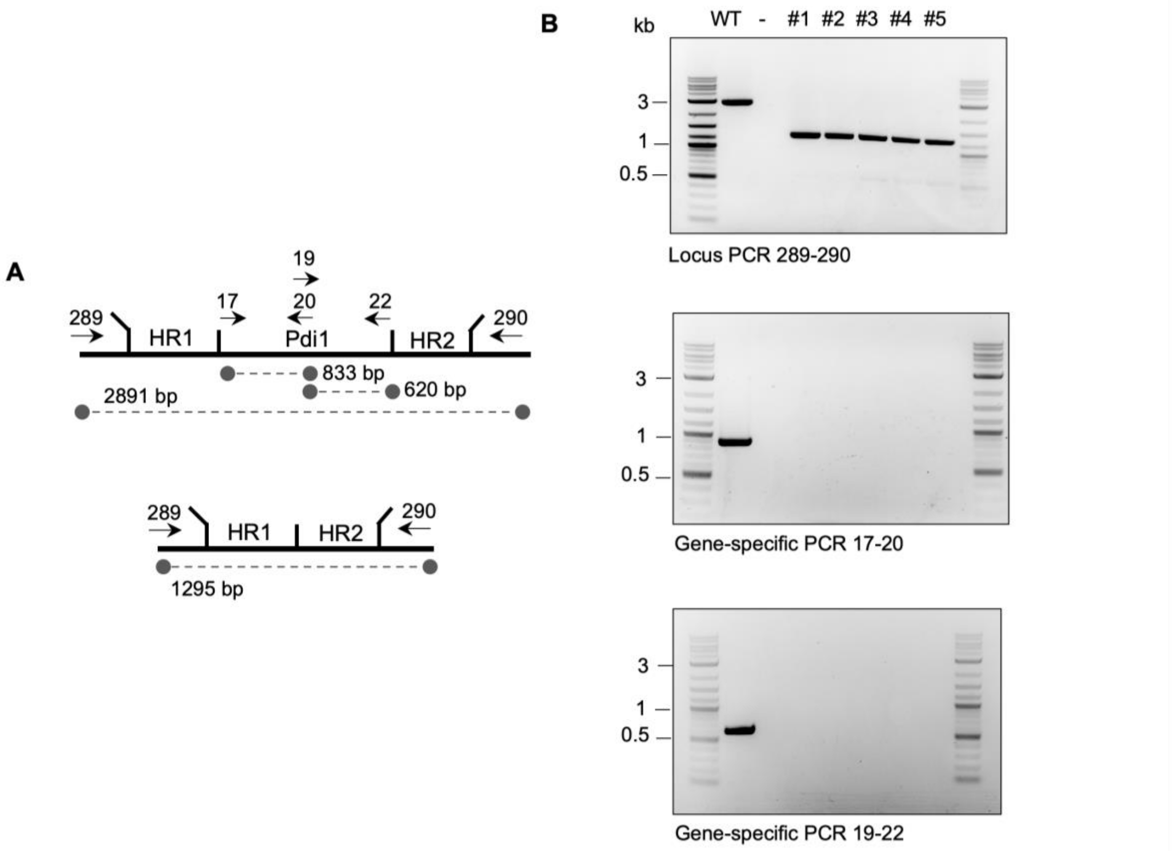

**Table S1**. List of plasmids used in this study.

| Plasmid # | Description | Resistance |
|---|---|---|
| pAR144 | BB1_1,2_repair template Pdi1 | KanaR |
| pAR147 | BB3cN_pGAP_23*_Pdi1_sg+_pLAT1_Cas9 | NTC |
| pAR148 | BB3cN_pGAP_23__Pdi1_sg-_pLAT1_Cas9 | NTC |
| pAR12 | BB3_AOXtt_ZeoR_pGAP_SsTrp_cycTT_Loxp | ZeoR |
| pAR107 | BB3_AOXtt_ZeoR_pGAP_scFvM_cycTT_Loxp | ZeoR |

**Table S2**. List of oligos used in this study. Golden Gate fusion sites are underlined in the sequence.

| Oligo name | Sequence | Use |
|---|---|---|
| AR279_Pdi_up_GG_F | GATCTAGGTCTCAGGAGATATGGTCCAGGAAAAAAACAAC | Homology repair template |
| AR280_Pdi_up_GG_R | ATATAGAAACgagaccGTACGTGTTCAATTATTGAAG | Homology repair template |
| AR281_Pdi_dw_GG_F | ACGTACggtctcGTTTCTATATTTACTGTAACTAGG | Homology repair template |
| AR282_Pdi_dw_GG_R | GTCATTGGTCTCTCATGTAGGCTGATGCTGTTGACA | Homology repair template |
| AR285_Pdi_up_F | TATGGTCCAGGAAAAAAACAAC | Homology repair template |
| AR286_Pdi_down_R | TAGGCTGATGCTGTTGACAC | Homology repair template |
| AR289_Pdi_ko_F | GAGACTCACGATCCTCGC | KO verification |
| AR290_Pdi_ko_R | GGACCCTGATGGGTATCTTC | KO verification |
| AR295_Pdi_sg_plus_F | atggtctcCCATGTTATAGCTGATGAGTCCGTGAGGACGAAAC GAGTAAGCTCGTCCTAT | Single guide |
| AR296_Pdi_sg_plus_R | AAACGAGTAAGCTCGTCCTATAAGAATGCAATTCAACgttttag agctagaaatagcaag | Single guide |
| AR297_Pdi_sg_minus_F | atggtctcCCATGTCCTATCTGATGAGTCCGTGAGGACGAAAC GAGTAAGCTCGTCATAG | Single guide |
| AR298_Pdi_sg_minus_R | AAACGAGTAAGCTCGTCATAGGAAGAAGAAGAAGGTTgtttta gagctagaaatagcaag | Single guide |
| qAR17_Pdi1_up_F | CCTCACGTTTTGGCAGAGTT | qPCR/KO verification |
| qAR19_Pdi1_mid_F | TCGGTAAGCATGCCAAGAAC | KO verification |
| qAR20_Pdi1_mid_R | TGTTCGTCAATTCTTGGTCTTGA | qPCR/ KO verification |
| qAR22_Pdi1_dw_R | GTCTGCCTCACTTTCAGCTT | KO verification |
| qAR5_HAC1_fw | GCGGCCCATGCTTCCAGAGAG | qPCR |
| qAR6_HAC1_bw | CGGTACCACCTAAGGCTTCCAACC | qPCR |
| qAR15_Kar2_F | TCTTGGCTGACTTTGGCGGCAT | qPCR |
| qAR16_Kar2_R | CCCGACTTCATCACACCGACACA | qPCR |
| qAR23_ERp38_up_F | TGTCCTTGGGTGTGAACAATTT | qPCR |
| qAR24_ERp38_up_R | GATTGCCGCCATTGATGCTA | qPCR |
| PpACT1_Up | CCTGAGGCTTTGTTCCACCCATCT | qPCR (internal standard) |
| PpACT1_Low | GGAACATAGTAGTACCACCGGACATAACGA | qPCR (internal standard) |

**Table S3.** Target protein sequences. Alpha-mating factor leader peptide is underlined at the beginning of the sequence.

| | |
|---|---|
| scFvM | MRFPSIFTAVLFAASSALAAPVNTTTEDETAQIPAEAVIGYSDLEGDFDVAVLPFSNSTNNGLL<br>FINTTIASIAAKEEGVSLEKRQEQLVESGGGLVQPGGSLRLSCAASGFDFSSHWIYWVRQAPG<br>KGLEWVSTIYTGSDSTYYATWAKGRFTISKDNSKNTVYLQMNSLRAEDTAVYYCARDLGGSS<br>STSYISDLWGQGTLVTVSSGGGGSGGGGSGGGGSELVLTQSPATLSLSPGERATLSCTLSS<br>AHKTYSIAWYQQKPGQAPRYLIQLKSDGSYTKGTGVPARFSGSSSGADRTLTISSLEPEDFAV<br>YYCSTDYATGYYVFGQGTKVEIKRGGSHHHHHH |
| Porcine Trypsinogen | MRFPSIFTAVLFAASSALAAPVNTTTEDETAQIPAEAVIGYSDLEGDFDVAVLPFSNSTNNGLL<br>FINTTIASIAAKEEGVSLDKREAEAFTDDDDKIVGGYTCAANSIPYQVSLNSGSHFCGGSLINSQ<br>WVVSAAHCYKSRIQVRLGEHNIDVLEGNEQFINAAKIITHPNFNGNTLDNDIMLIKLSSPATLNS<br>RVATVSLPRSCAAAGTECLISGWGNTKSSGSSYPSLLQCLKAPVLSDSSCKSSYPGQITGNMI<br>CVGFLEGGKDSCQGDSGGPVVCNGQLQGIVSWGYGCAQKNKPGVYTKVCNYVNWIQQTIA<br>AN |
